# A novel phylogenetic analysis and machine learning predict pathogenicity of human mtDNA variants

**DOI:** 10.1101/2020.01.10.902239

**Authors:** Bala Anı Akpınar, Paul O. Carlson, Ville O. Paavilainen, Cory D. Dunn

## Abstract

Linking mitochondrial DNA (mtDNA) variation to clinical outcomes remains a formidable challenge. Diagnosis of mitochondrial disease is hampered by the multicopy nature and potential heteroplasmy of the mitochondrial genome, differential distribution of mutant mtDNAs among various tissues, genetic interactions among alleles, and environmental effects. Here, we describe a new approach to the assessment of which mtDNA variants may be pathogenic. Our method takes advantage of site-specific conservation and variant acceptability metrics that minimize previous classification limitations. Using our novel features, we deploy machine learning to predict the pathogenicity of thousands of human mtDNA variants. Our work demonstrates that a substantial fraction of mtDNA changes not yet characterized as harmful are, in fact, likely to be deleterious. Our findings will be of direct relevance to those at risk of mitochondria-associated metabolic disease.

## INTRODUCTION

Because of the critical roles that mitochondria play in metabolism and bioenergetics, mutation of mitochondria-localized proteins and ribonucleic acids can adversely affect human health (Alston *et al*, 2017; Suomalainen & Battersby, 2018; Khan *et al*, 2020; Russell *et al*, 2020). Indeed, at least one in 5000 people (Gorman *et al*, 2015) is estimated to be overtly affected by mitochondrial disease. While a very limited number of mitochondrial DNA (mtDNA) lesions can be directly linked to human illness, the clinical outcome for many other mtDNA changes remains ambiguous (Vento & Pappa, 2013). Heteroplasmy among the hundreds of mitochondrial DNA (mtDNA) molecules found within a cell (Stewart & Chinnery, 2015; Hahn & Zuryn, 2019; Wei & Chinnery, 2020), differential distribution of disease-causing mtDNA among tissues (Boulet *et al*, 1992), and modifier alleles within the mitochondrial genome (Wei *et al*, 2017; Elliott *et al*, 2008) magnify the difficulty of interpreting different mtDNA alterations. Mito-nuclear interactions and environmental effects may also determine the outcome of mitochondrial DNA mutations (Wolff *et al*, 2014; Hill *et al*, 2019; Matilainen *et al*, 2017; Turnbull *et al*, 2018). Beyond the obvious importance of resolving the genetic etiology of symptoms presented in a clinical setting, the rapidly increasing prominence of direct-to-consumer genetic testing (Phillips *et al*, 2018) calls for an improved understanding of which mtDNA polymorphisms might affect human health (Blell & Hunter, 2019).

Simple tabulation of mtDNA variants found among healthy or sick individuals (Whiffin *et al*, 2017) may be of limited utility in predicting how harmful a variant may be. Differing, strand-specific mutational propensities for mtDNA nucleotides at different locations within the molecule (Tanaka & Ozawa, 1994; Faith & Pollock, 2003; Reyes *et al*, 1998) should be taken into account when assessing population-wide data, yet allele frequencies are rarely, if ever, normalized in this way. Population sampling biases and recent population bottleneck effects can lead to misinterpretation of variant frequencies (Zuk *et al*, 2014; Chheda *et al*, 2017; Keinan & Clark, 2012; Landry *et al*, 2018; Pirastu *et al*, 2020). Mildly deleterious variants arising in a population are slow to be removed by selection (Nachman, 1998; Nachman *et al*, 1996), leading to a false prediction of variant benignancy. Finally, a lack of selection against variants that might act in a deleterious manner at the post-reproductive stage of life also makes likely the possibility that some mtDNA changes will contribute to age-related phenotypes while avoiding overt association with mitochondrial disease (Maklakov *et al*, 2015; Medawar, 1952; Cui *et al*, 2019; Williams, 1957; Wallace, 1994).

Examining evolutionary conservation by use of multiple sequence alignments offers important assistance when predicting a variant’s potential pathogenicity (Raychaudhuri, 2011; Tang & Thomas, 2016a). However, caveats are also associated with predicting mutation outcome by the use of these alignments. First, while knowledge of amino acid physico-chemical properties is widely considered to be informative regarding whether an amino acid substitution may or may not have a damaging effect on protein function (Dayhoff *et al*, 1978), the site-specific acceptability of a given substitution is ultimately decided within the context of its local protein environment (Zuckerkandl & Pauling, 1965). Second, sampling biases and improper clade selection may lead to inaccurate clinical interpretations regarding the relative acceptability of specific variants (Zuk *et al*, 2014; Chheda *et al*, 2017; Keinan & Clark, 2012; Landry *et al*, 2018). Third, alignment (Kawrykow *et al*, 2012; Iantorno *et al*, 2014) and sequencing errors (Chen *et al*, 2017; Smith, 2019) may falsely indicate the acceptability of a particular mtDNA substitution.

Here, we have deployed a methodology to calculate, by a novel analysis of available mammalian genomes, the relative conservation of human mtDNA-encoded positions. Moreover, we infer ancestral direct substitutions within mammals and test whether they match substitutions from the human reference sequence, providing further knowledge regarding the potential pathogenicity of any human mtDNA substitution. By subsequent application of machine learning, we demonstrate that a surprising number of uncharacterized mtDNA mutations carried by humans are likely to promote disease. We provide our predictions, which should be of great utility to clinicians and to those studying mitochondrial disease.

## RESULTS

### Mapping apparent substitutions to a phylogenetic tree allows calculation of relative positional conservation in mtDNA-encoded proteins and RNAs

We previously developed an empirical method for detection and quantification of mtDNA substitutions mapped to the edges of a phylogenetic tree (Dunn *et al*, 2020). Here, we have extended our approach toward prediction of human mitochondrial variant pathogenicity. First, we retrieved full mammalian mtDNA sequences from the National Center for Biotechnology Information Reference Sequence (NCBI RefSeq) database and extracted each RNA or protein-coding gene using the *Homo sapiens* reference mtDNA as a guide. Next, we aligned the resulting protein, tRNA, and rRNA sequences, concatenated the sequences of each species based upon molecule class, and generated phylogenetic trees using a maximum likelihood approach. Following tree generation, we performed ancestral prediction to reconstruct the character values of each position at every bifurcating node. Using the sequences of extant species and the predicted ancestral node values, we subsequently analyzed each edge of the tree for the presence or absence of substitutions at each aligned human position. We subsequently sum all substitutions at a given position that occur along all tree edges to generate a new metric, the total <u>s</u>ubstitution <u>s</u>core (TSS, Figure 1A). The TSS should surpass metrics that consider positional character frequencies derived from multiple sequence alignments as a proxy of conservation, as character frequencies are highly sensitive to sampling biases among input sequences. Moreover, many site-specific measurements of variability, such as Shannon entropy, are limited in dynamic range and benefit minimally from the rapid increase in available genomic information. In contrast, the dynamic range of the TSS is very wide, and potentially unlimited, continuously benefitting from the accretion of new sequence information.

**Figure 1:**
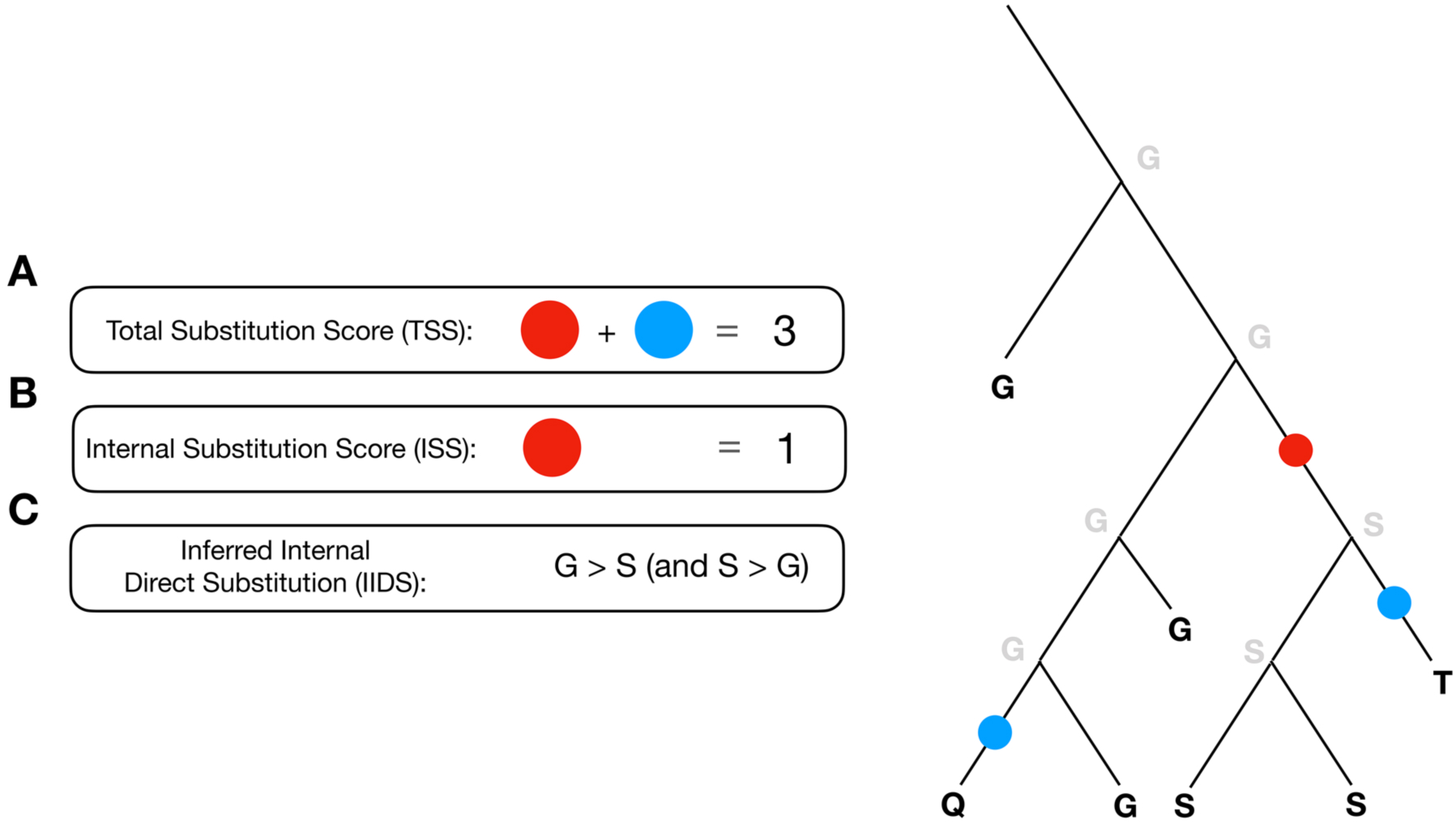
Calculation of Total Substitution Scores, Internal Substitution Scores, and Inferred Internal Direct Substitutions. Phylogenetic trees are produced for each macromolecule class (protein, rRNA, tRNA), then ancestral values were predicted for each node of the tree. Glycine is ancestral in this simplified example, with substitution to serine, threonine, and glutamine at internal or external branches. (A) Next, substitutions at an alignment position that occur at tree edges are summed to produce the total substitution score (TSS). (B) To minimize tabulation of artifactual changes related to alignment and sequencing errors, only substitutions at internal edges are summed to generate the internal substitution score (ISS). (C) Using the predicted node values, inferred internal direct substitutions (IIDSs) along any internal edge are determined, preferentially taking into account epistasis involving the site of interest and avoiding the effects of sequencing and alignment errors.

Furthermore, by excluding edges from analysis that lead directly to extant sequences, one can further minimize effects of alignment errors and sequencing errors that may lead to eventual misinterpretation of variant pathogenicity. Moreover, mutations mapped to internal edges are more likely to represent fixed changes informative for the purposes of disease prediction, while polymorphisms that have not yet been subject to selection of sufficient strength or duration might be expected to complicate predictions of variant pathogenicity (Nachman *et al*, 1996; Nachman, 1998). Summation of substitutions only at these internal edges provides an internal <u>s</u>ubstitution <u>s</u>core (ISS, Figure 1B).

When calculated for protein and RNA sites encoded by mammalian mtDNA, it is clear that the TSS (and the ISS, not shown) provides an excellent readout of relative conservation at, and consequent functional importance of, each alignment position. When comparing TSS data from different mtDNA-encoded proteins, our findings are consistent with previous results, obtained by alternative methodologies, demonstrating that the core, mtDNA-encoded subunits of Complexes III and IV tend to be the most conserved, while positions within the mtDNA-encoded polypeptides of Complex I and Complex V tend to be less well conserved (da Fonseca *et al*, 2008; Nabholz *et al*, 2013) (Figure 2A). Examination of the structures of these complexes indicate that, indeed, the most conserved residues are preferentially localized near the key catalytic regions of each complex (not shown). Within each protein, there was, as expected, a spectrum of site conservation values, also illustrated by plotting a distribution of TSS values across each polypeptide (Figure S1). Nearly all analyzed protein positions appeared to be under some selective pressure and are not saturated with mutations, with TSS values existing far from the maximal values that can be achieved within this phylogenetic analysis of mammals. Selective pressure on most aligned sites is also observed when examining mtDNA-encoded tRNAs and rRNAs (Figure 2B and Figure S2).

**Figure 2:**
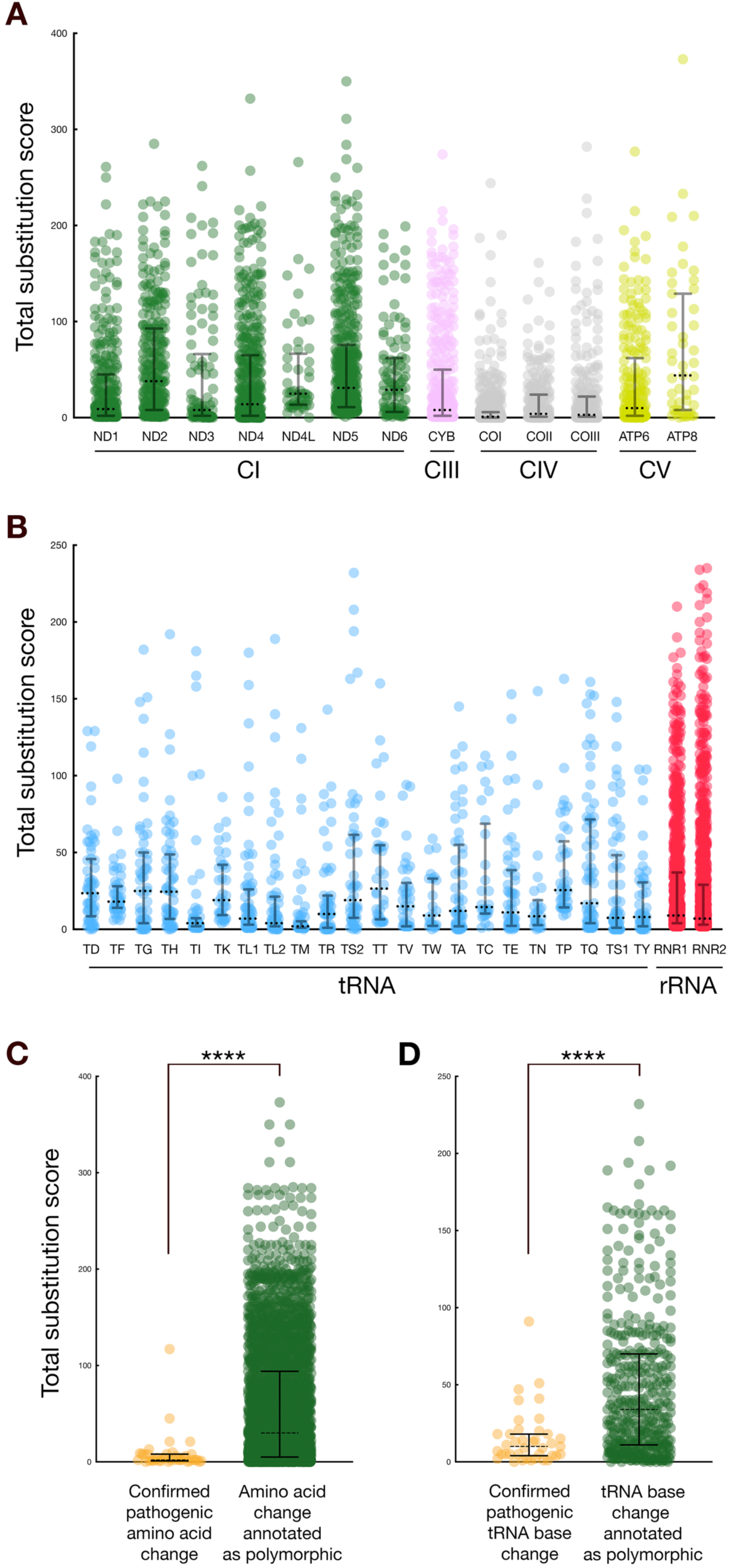
Most well-aligned positions encoded by the human mitochondrial genome are under selective constraint. (A) Nearly all amino acids in mammalian mtDNA-encoded proteins are under selection. The TSS analysis of alignment sites within mtDNA-encoded proteins, clustered by complex (89% of total amino acid positions, gapped 1.5% or less), are provided. (B) Nearly all ribonucleotides across mammalian mtDNA-encoded tRNAs and rRNAs are under selection. The TSS analysis of alignment sites within mtDNA-encoded tRNAs (75% of total tRNA positions, gapped 1.5% or less) or rRNAs (77% of total tRNA positions, gapped 1.5% or less) are provided. (C-D) Confirmed pathogenic mutations in protein and tRNAs are associated with low total substitution scores. TSSs for positions of variants annotated in the MITOMAP database as confirmed pathogenic substitutions or as polymorphisms are plotted for (C) protein positions analyzed using mammalian data and (D) tRNA positions analyzed using mammalian data. All analyzed positions are gapped at equal to or less than 1.5%. ****, Kolmogorov-Smirnov test approximate P-value of <0.0001. Dotted horizontal lines, median, with solid horizontal lines providing the interquartile range.

Beyond summation of substitutions across a phylogenetic tree, the inferred ancestral and descendent characters at each edge of the phylogenetic tree can also be examined following generation of the substitution map and can provide important information regarding what changes to mtDNA-encoded macromolecules might be deleterious or not. Specifically, if an inferred direct substitution from the human reference character to the mutant character (or the inverse, assuming the time-reversibility of character substitutions) is predicted along the edge of a phylogenetic tree, then such a change at a given position might be expected to be less deleterious than an inferred direct substitution to or from the human character that was never encountered over the evolutionary history of a clade. In contrast, the simple presence or absence of a character at an alignment position, without the context of its ancestral character, will fail to reflect epistatic relationships that involve the position of interest (Kimura, 1985; Kondrashov *et al*, 2002; Marini *et al*, 2010). A further focus upon inferred direct substitutions occurring at internal tree edges, which we call internal inferred <u>d</u>irect <u>s</u>ubstitutions (IIDSs) (Figure 1C), avoids the confounding effects of sequencing and alignment errors (Chen *et al*, 2017; Smith, 2019) and of the abovementioned, incomplete selection against newly arisen, mildly deleterious alleles (Nachman *et al*, 1996; Nachman, 1998).

### Substitution scores and inferred direct substitutions can be linked to human mtDNA variant pathogenicity

Since summation of detected substitutions across a phylogenetic tree provides a robust measure of relative conservation at different macromolecular positions, we were confident that a phylogenetic analysis that includes TSSs would also provide information about the pathogenicity of human mtDNA variants. To test this possibility, we focused our attention upon harmful substitutions annotated within the MITOMAP database of pathogenic mtDNA alterations (Lott *et al*, 2013). Indeed, we detected a clear relationship between confirmed pathogenicity and conservation at mtDNA-encoded amino acid positions, as reflected by positional TSSs (Figure 2C). Similarly, there was a strong link between tRNA mutation pathogenicity and TSSs obtained from mammalian tRNA data (Figure 2D). The paucity of confirmed pathogenic mitochondrial rRNA mutations in the MITOMAP database made comparisons using this class of molecules impractical, yet we expect future confirmation of additional pathogenic mutations in mitochondria-encoded rRNAs to permit further analyses. Together, findings obtained by phylogenetic analysis of mitochondria-encoded proteins and tRNAs indicate that TSSs are a valuable asset when predicting which mtDNA mutations might lead to disease.

To test the possibility that IIDSs, like TSSs, may predict pathogenicity, we compared human variants confirmed to cause disease to those variants currently labelled as polymorphisms within the MITOMAP database. Pathogenic human amino acid changes from the reference sequence are rarely encountered among the internal edges of a mammalian phylogenetic tree (Figure 3A). In contrast, reportedly polymorphic alleles were much more likely to have the amino acid substitution identifiable along internal edges of the mammalian tree. For nucleotide substitutions in tRNAs, a similar analysis comparing confirmed pathogenic variants and variants annotated as polymorphisms detected no statistically significant relationship between pathogenicity and the detection of IIDSs.

**Figure 3:**
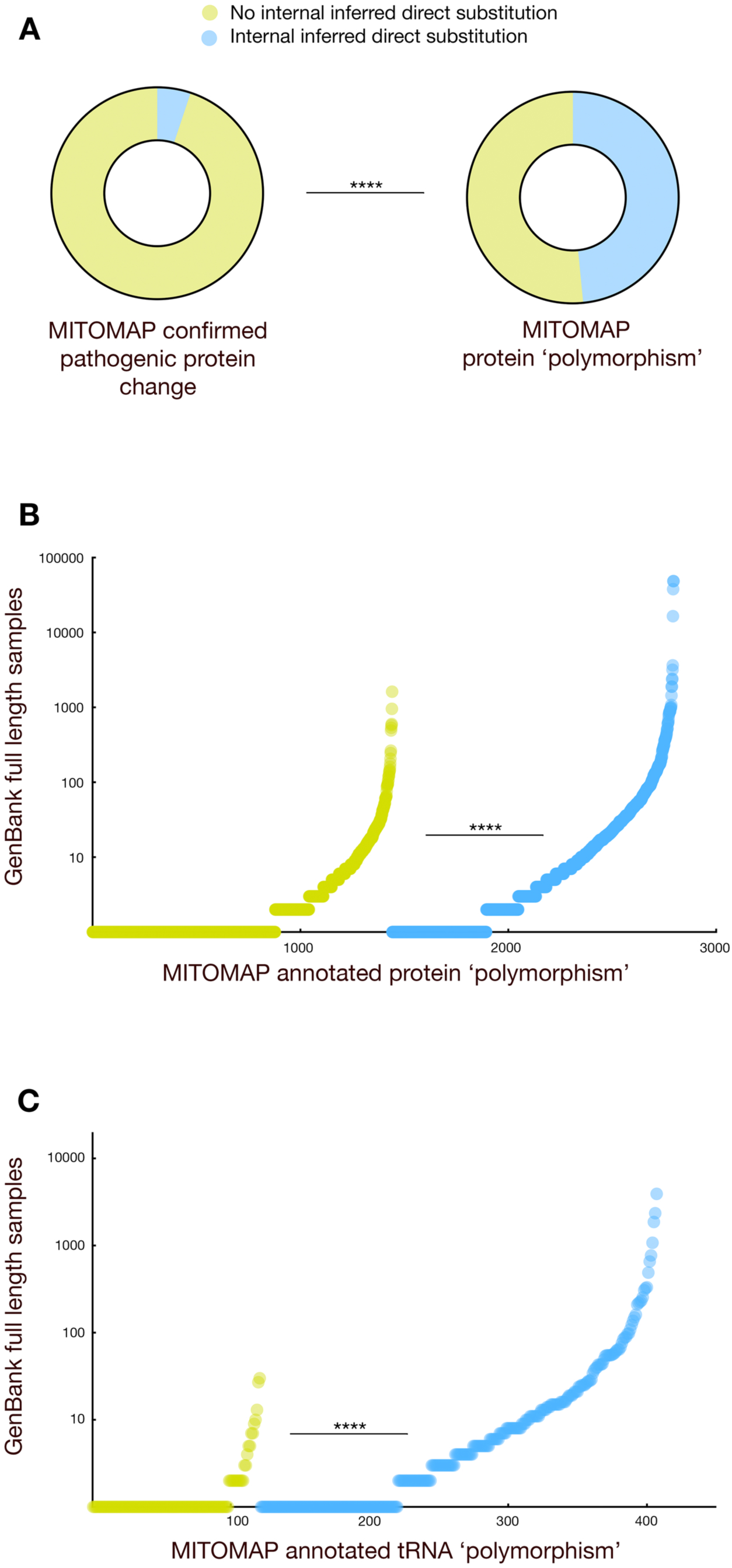
Analysis of inferred internal direct substitutions indicates that variants currently annotated as polymorphic may be deleterious. (A) Inferred internal direct substitutions within the mammalian phylogenetic tree correspond to a reduced likelihood of confirmed pathogenicity for mitochondrial protein coding variants. Confirmed pathogenic protein substitutions and substitutions currently annotated as polymorphisms, both retrieved from MITOMAP, were tested for the presence of an IIDS. All analyzed positions are gapped at equal to or less than 1.5%. ****, Fisher’s two-sided exact test P-value of <0.0001. (B) Population frequency distributions differ when amino acid substitutions from the human reference, which are currently annotated as polymorphic in MITOMAP, are classified based upon the presence or absence of an IIDS. All analyzed positions are gapped at equal to or less than 1.5%. (C) as in (8), but for tRNA substitutions. ****, Kolmogorov-Smirnov test approximate P-value of <0.0001.

### Widespread, uncharacterized pathogenicity among human mtDNA polymorphisms

A subset of substitutions currently annotated as ‘polymorphic’ in the MITOMAP database were distinguished by very low TSS scores at the relevant alignment position, suggesting that mutations at those positions could, in fact, be deleterious. Moreover, when considering polymorphisms, the absence of IIDSs within the mammalian phylogenetic tree that would match particular human variations suggested that additional pathogenicity may exist among human mtDNA substitutions currently considered polymorphic. In order to explore whether additional pathogenic mutations might indeed be discovered among variants currently tabulated as ‘polymorphic,’ we tested whether protein polymorphisms tabulated in MITOMAP that cannot be directly linked to the human reference character along internal edges of the mammalian phylogenetic tree are under purifying selection. Selection was assessed by determining the number of full-length mtDNA samples found in GenBank (Benson *et al*, 2013) that were assigned within MITOMAP to a given variant. Measuring selection by assessment of population frequencies can be problematic due to potential sampling biases and population bottlenecks (Auer & Lettre, 2015), timing of mutation arrival within an expanding population (Luria & Delbrück, 1943; Rosche & Foster, 2000), and the divergent nucleotide- and strand-specific mutational propensities of mtDNA (Tanaka & Ozawa, 1994; Faith & Pollock, 2003; Reyes *et al*, 1998). Even so, the distribution of variant frequencies among full-length sequences in GenBank was strikingly different for those mutations for which an IIDS could be identified in our mammalian trees of proteins (Figure 3B), and even tRNAs (Figure 3C), when compared to those for which an IIDS could not be identified. These results indicate that many human mtDNA protein and tRNA variants currently considered to be polymorphic are, in fact, harmful, and these findings further validate the development of the IIDS as a feature useful for determining variant pathogenicity.

### A support vector machine predicts harmful mtDNA variants

Given the clear presence of deleterious substitutions among so far uncharacterized variants, we sought a high-throughput method that could, with confidence, identify these potentially deleterious substitutions. Toward this goal, we turned to machine learning. Specifically, we deployed a support vector machine (SVM) (Cortes & Vapnik, 1995), designed to minimize risk of misclassification and able to take advantage of non-linear features (Bhavsar & Ganatra, 2012), to predict the risk of mtDNA variants using our conservation features. Positive training sets for our SVM consisted of protein or tRNA mutations annotated as confirmed pathogenic alterations within the MITOMAP database (Lott *et al*, 2013). Negative training sets included variants annotated as polymorphic in MITOMAP and also characterized by the highest counts of homoplasmic alleles within HelixMTdb, a database emerging from a survey of nearly 200,000 human samples (Bolze *et al*, 2019). Beyond these training variants, our total set of variants for prediction included substitutions tabulated in MITOMAP, substitutions indexed in HelixMTdb, and simulated substitutions from the reference sequence not yet encountered in the clinic. Analyzed positions were gapped within less than or equal to 1.5% of analyzed sequences. As SVM features, we exploited TSS, ISS, and IIDS. However, since SVMs may beneficially incorporate features, even with theoretical shortcomings, in unanticipated ways, we also expanded our feature set to include other features related to conservation. These additional features included the number of characters found at alignment positions (internal edges and all edges), whether the mutant character is ever found at internal nodes, and positional Shannon entropy.

For variants found within protein-coding genes, our SVM approach clearly permitted a robust and global prediction of which polymorphisms not yet classified as pathogenic may cause or contribute to disease (Figure 4A and Table S1). Importantly, when examining the behavior of protein-coding variants within our negative training sets, only one false-positive was obtained. Furthermore, few confirmed pathogenic mutations were found among the predicted negative set. The proficiency of our model in correctly identifying true positives in the training set without a corresponding loss in specificity is highlighted by the associated Receiver Operating Characteristic (ROC) curve [area under the receivership curve (AUC), 0.96] (Figure S3A). The Matthews Correlation Coefficient (MCC), reflecting the overall performance of our classifier (Guilford, 1954; Chicco & Jurman, 2020), was 0.89, indicating a model with high proficiency at separating our positive and negative training sets. Accuracy for our model was calculated to be 0.95, precision was 0.97, sensitivity was 0.90, specificity was 0.98, and our F-score was 0.94, all of which are outcomes that also suggest outstanding classifier performance on variation within mitochondria-encoded polypeptides. Of features assessed by our SVM, all of which are related to site conservation among mammals, IIDS stood out as dominant in providing predictive value (Figure S3B).

**Figure 4:**
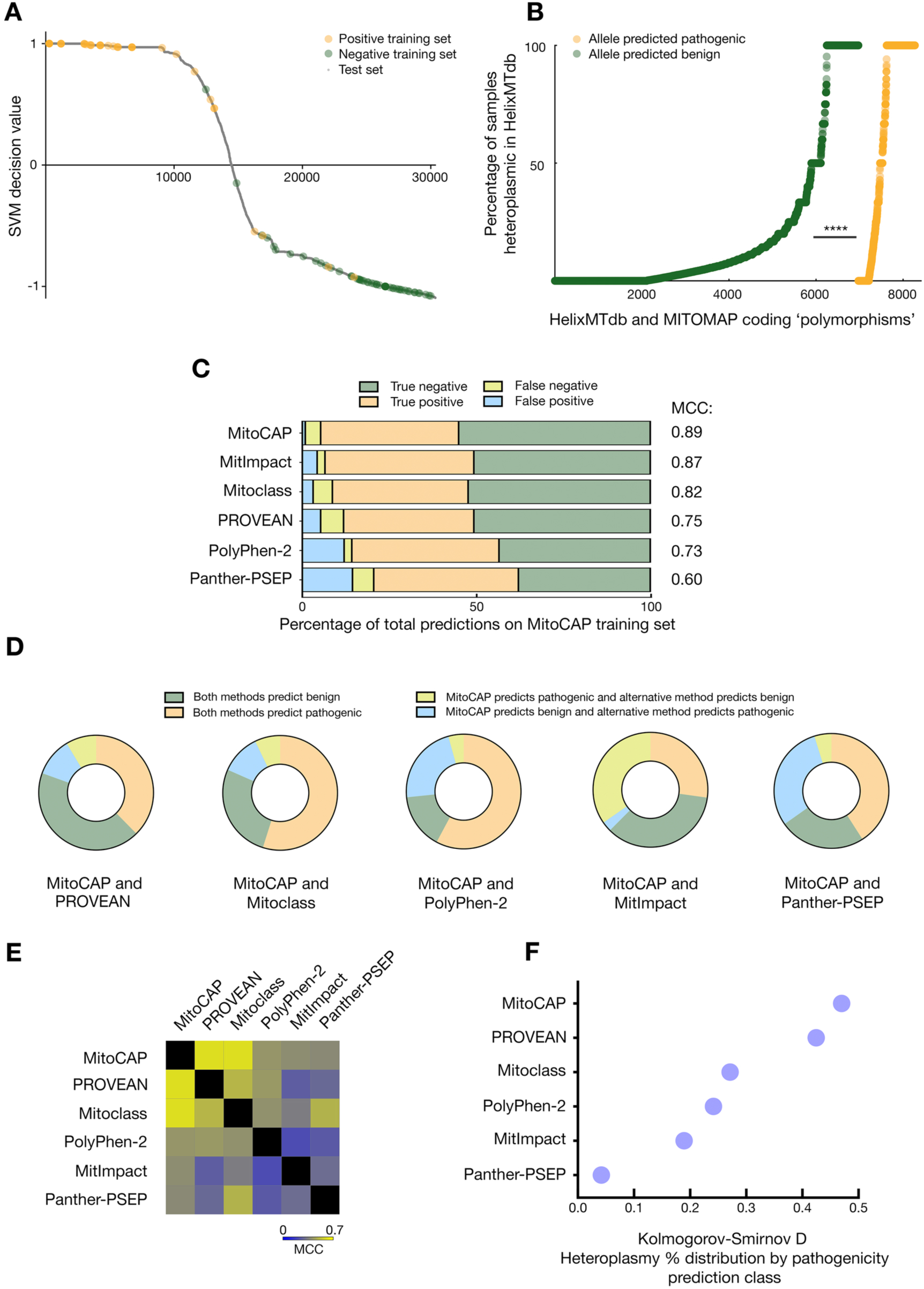
The MitoCAP classifier can successfully predict the pathogenicity of protein-coding mtDNA variants currently annotated as polymorphisms. (A) SVM-based prediction of pathogenicity for protein-coding variants effectively separates the positive and negative training sets, which are superimposed on the curve of variants and decision values. Variants are arrayed by decreasing SVM decision values on the x-axis. (B) MitoCAP classification of protein variants leads to distinct distributions of heteroplasmy propensity. The fractions of heteroplasmic HelixMTdb samples for a given variant are plotted in order of increasing value on the x-axis and separated by MitoCAP pathogenicity prediction. Plotted variants are annotated as ‘polymorphic’in MITOMAP or are unannotated and do not belong to the positive or negative training sets. ****, Kolmogorov-Smirnov test approximate P-value of <0.0001. (C) Comparison of protein substitution classifier behavior using the MitoCAP training set. The percentage of total predictions on the MitoCAP training set that are true negative, true positive, false negative, or false positive are reflected, as well as the calculated MCC. (D-E) MitoCAP predictions on substitutions in protein-coding genes are most similar to those of the PROVEAN classifier. (D) The fraction and characterization of instances where both approaches are in agreement or are discordant is represented. Both classifiers had to have made predictions in order for the comparison to enter the analysis, and all amino acid substitutions, including those of training sets, are included. (E) Comparison of prediction outcomes between each pair of methods, including all test and training set variants, was performed using MCC calculations and represented as a heat map. (F) MitoCAP classification of substitutions in mitochondrial proteins best differentiates heteroplasmy frequency distributions of unannotated variants. The Kolmogorov-Smirnov D statistic for each pair of classification-based distributions [(B) and Figure S5] is shown. The associated approximate P-values associated with each test are <0.0001, with the exception of Panther-PSEP (P value <0.05).

To further validate our approach, we turned to the thousands of uncharacterized mtDNA changes encountered during generation of the HelixMTdb. Data presented above (Figure 3B and 3C) and previous studies (Ye *et al*, 2014; Elliott *et al*, 2008; Wei & Chinnery, 2020) support the idea that pathogenic alleles reside among the set of uncharacterized changes and mutations currently annotated as polymorphic. Moreover, frequent clinical encounters with a variant observed in a heteroplasmic state (not consisting of 100% of the molecules in a cell) can often be a signal of pathogenicity (Stewart & Chinnery, 2015; Hahn & Zuryn, 2019). Therefore, we reasoned that by first separating unannotated variants by SVM prediction class, then comparing the fraction of instances that each allele within that class was determined to be heteroplasmic within an individual, we might assess our prediction capability. We note that heteroplasmy was not a feature used for development of our prediction model. Upon analysis, heteroplasmy profiles were quite clearly distinct (Figure 4B) when comparing those variants that we predict to be harmful and those variants that we predict to be benign, and we obtained a Kolmogorov-Smirnov D statistic of 0.471, suggesting that our methodology has quite successfully revealed alleles in protein-coding genes that are likely to cause metabolic disease. We have named our approach MitoCAP, for “<u>Mito</u>chondrial Disease Predictions Using <u>C</u>hanges <u>A</u>cross <u>P</u>hylogeny”.

Other classifiers provide their own predictions, or otherwise provide the opportunity to predict, whether some or all of the potential changes in mtDNA-encoded proteins may be deleterious. We selected several of these methods for comparison with MitoCAP. We focused our attention primarily upon two recently developed and mitochondria-specific classifiers: Mitoclass (Martín-Navarro *et al*, 2017) and the meta-classifier MitImpact (Castellana *et al*, 2015). We also included two other tools that allow batch submission of substitution input, PolyPhen-2 (Adzhubei *et al*, 2013) and PROVEAN (Choi *et al*, 2012), promoting further evaluation of mitochondria-specific classifiers. Finally, we chose Panther-PSEP (Tang & Thomas, 2016b) which, like MitoCAP, takes advantage of phylogenetic tree information while making predictions. These different classifiers do include some convergent concepts within their prediction schemes. However, we expected that our use of novel phylogenetic and conservation features would lead to tangible improvement in the prediction of mtDNA variant pathogenicity.

First, we tested how the compared methods would fare against the training set that was used to build our own prediction model for variants in protein-coding genes. It may be challenging, in an unbiased manner, to compare metrics of prediction success that we obtained using our own training sets to those achieved by other methods using our training sets. However, selection of variants for inclusion within our training sets was based upon conservative measures of pathogenicity or benignancy, and not upon our own interpretation of individual variant hazard. Therefore, we should expect other classifiers to also perform well on our training set. Of the six methods compared, the two methods specifically designed to assess variants within mitochondrial protein coding genes, MitImpact and Mitoclass, performed second and third best on our training set, as reflected by MCC values. These classifiers were followed by PROVEAN, PolyPhen-2, and Panther-PSEP (Figure 4C). MitoCAP also scored best against our training set when considering most auxiliary measures of prediction proficiency (Figure S4).

We further explored the magnitude by which the underlying prediction models might differ from one another. To further investigate this possibility, we first plotted the level of agreement between MitoCAP other methods when assessing all classified variants, and we noted a pronounced lack of overlap between our MitoCAP predictions and the predictions of other methods (Figure 4D). For example, MitImpact appears to predict that a substantial number of alleles that we predict as pathogenic are actually innocuous. In contrast, Panther-PSEP predicts that a pronounced number of alleles that we call benign are actually deleterious.

In order to more quantitatively assess how similar each pair of prediction outcomes may be to one another, we next applied MCC calculations as a way to measure prediction correspondence. During these analyses (Figure 4E), the maximum MCC value calculated during comparisons between methods was 0.61, obtained during comparison of MitoCAP and PROVEAN, and a similar score of 0.60 was the outcome of comparing the MitoCAP and Mitoclass predictors. All other comparisons provided MCC values of 0.5 or (sometimes substantially) less. Therefore, MitoCAP seems to provide predictions for unannotated variants that are most in accordance with, but not identical to, those produced by PROVEAN and Mitoclass.

But which of these prediction approaches is most likely to appropriately characterize unannotated variants? When heteroplasmy data for unannotated variants in HelixMTdb are analyzed for other prediction methods (Figure S5), as performed above for MitoCAP, MitoCAP best separated variants into classes with different heteroplasmy propensities and achieved the highest Kolmogorov-Smirnov D score (Figure 4F). Consequently, our prediction method appears to be the most successful among the tested classifiers in identifying harmful mtDNA variants within protein-coding genes.

For mitochondria-encoded tRNAs, notable separation between positive and negative training sets (Figure S6A and Table S2) was also achieved by MitoCAP. When considering our tRNA training set, the MitoCAP MCC value was 0.82, accuracy was 0.91, precision was 0.86, sensitivity was 0.95, specificity was 0.88, and the F-score was 0.90. The strong performance of our classifier is exemplified by the ROC curve (Figure S3C) and the associated AUC of 0.94. The number of characters seen at internal nodes for positions of interest, as well as the TSS and ISS, appeared to play the strongest role during development of the SVM prediction model (Figure S3D). When samples outside of our training set from our two predictive classes were plotted for heteroplasmy classification frequency in HelixMTdb, we again noted a significant difference in the two distributions (Figure S6B), indicating successful prediction of previously unanticipated pathogenicity. When compared to two other methods of tRNA variant pathogenicity prediction, PON-mt-tRNA (Niroula & Vihinen, 2016) and MitoTIP (Sonney *et al*, 2017), the MitoTIP classifier performed better when considering the MitoCAP training set (Figure S6C and Figure S7). All three methods tested appear to diverge substantially from one another in their pathogenicity predictions (Figure S6D and Figure S6E). MitoTIP most distinctly separated distributions of heteroplasmy frequency within different predictive classes of unannotated variants (Figure S6F and Figure S8). However, superior separation of heteroplasmy distributions by MitoTIP may not be surprising, since MitoTIP incorporates whether variants have been encountered as heteroplasmic during its classifications, while MitoCAP does not.

Taken together, our analyses indicate that MitoCAP appears to be the most proficient among the compared methods in predicting pathogenicity of variants in mtDNA-encoded proteins, while alternative methods may outperform MitoCAP during classification of tRNA variants.

## DISCUSSION

We describe here a methodology that allows improved quantification of the relative conservation of sites within and between genes, RNAs, and proteins. TSS and ISS values provide a measure of conservation that can minimize errors in conservation calculation that result from sampling biases. Importantly, these scores are of theoretically unlimited dynamic range and will benefit from the continuous expansion of available sequence information. Even nearly identical sequences can be utilized by our approach, allowing for an ever-increasing input dataset that can be deployed toward calculation of site-specific conservation.

Beyond the development of a novel quantitative measure of conservation, we also use ancestral predictions to generate IIDSs - a binary read-out of site-specific substitution information. Researchers often consult generalized substitution matrices when predicting whether a change may be harmful or not (Dayhoff *et al*, 1978; Jones *et al*, 1992), yet amino acid exchangeability matrices change when moving across clades (Zou & Zhang, 2019), and successful substitution of any given character occurs in the context of a very specific local environment (Zuckerkandl & Pauling, 1965; Tang & Thomas, 2016a). By mapping substitutions to phylogenetic tree edges, ample sequence data can allow identification of IIDSs within the context of a particular macromolecular position, thereby improving prediction of variant pathogenicity. We note that focusing upon IIDSs, rather than the simple presence or absence of a character at a site, can indirectly integrate information about potential epistatic interactions that permit or block a substitution from being successfully established within a lineage.

Protein variants of confirmed pathogenicity were clearly characterized by low substitution scores and by an absence of IIDSs. But how should one weigh these and other available factors when classifying, in an unbiased and high-throughput manner, other variants already encountered or waiting to be discovered among human mtDNAs? To address this question, we deployed a machine learning approach, trained on existing data present in the MITOMAP database, toward determination of mtDNA mutation pathogenicity. Our apparent false-positive rate for protein coding variants appears to be extremely low, which is essential when avoiding the potential psychological effects of a false-positive test (Stewart-Brown & Farmer, 1997; Committee on Bioethics, 2001). Moreover, metrics of prediction capability for mitochondrial amino acid substitutions indicate very strong proficiency at pathogenicity prediction (MCC of 0.89), and specificity and sensitivity metrics resulting from our SVM predictions appear exemplary (AUC > 0.96). Importantly, comparison of multiple approaches by measuring the difference in heteroplasmy propensity, a common indicator of pathogenicity (Ye *et al*, 2014), within different predicted classes suggests that we can successfully outperform other methods specifically designed to classify mitochondrial variants, as well as several approaches focused more generally upon protein substitutions.

The MitoCAP predictions that we provide allow for improved comprehension of which mtDNA variants identified within a patient may be linked to mitochondrial disease. Our approach also allows researchers focusing upon the fundamental aspects of mitochondrial disease to rapidly prioritize variants identified in the clinic for directed study.

### mtDNA variants predicted to be pathogenic may lead to cryptic and age-related mitochondrial disease

Our results are congruent with earlier analyses suggesting that harmful mtDNA substitutions may be common within the human population (Ye *et al*, 2014; Elliott *et al*, 2008; Nachman *et al*, 1996). So why have so many of these deleterious changes not yet been classified as pathogenic within the clinic?

First, if a mutation is relatively common within the population, clinicians may inappropriately determine that the mtDNA variant is unlinked to patient symptoms (Whiffin *et al*, 2017). However, population frequency is not currently, at least when taken in isolation, a highly reliable predictor of clinical outcome, since population counts require substantial correction for strand- and nucleotide-specific mtDNA mutational biases (Tanaka & Ozawa, 1994; Faith & Pollock, 2003; Reyes *et al*, 1998), and sampling biases are typically a hazard when carrying out population-wide studies (Zuk *et al*, 2014; Chheda *et al*, 2017; Keinan & Clark, 2012; Landry *et al*, 2018).

Second, deleterious mtDNA variants often must rise beyond a certain threshold among the hundreds of mtDNA molecules potentially resident within a cell before overt symptoms can manifest (Stewart & Chinnery, 2015; Russell *et al*, 2020). Concordantly, our data suggest a strong propensity for heteroplasmy in the set of substitutions that we predict to be pathogenic, but are not yet clinically annotated as disease-associated. Differential distribution of mtDNA variants within a carrier, either established during development or resulting from bottleneck effects in renewable and non-mitotic tissues (Zhang *et al*, 2018; Stewart & Chinnery, 2015), may generate clones with a high proportion of deleterious mutations and may lead to complex, tissue-specific outcomes (Greaves *et al*, 2006; Nekhaeva *et al*, 2002; Bratic & Larsson, 2013; Fayet *et al*, 2002). Moreover, the phenomena described above may lead to age-related symptoms not easily classified as mitochondrial disease, since even relatively common mtDNA sequence variants have been suggested to contribute to diseases like diabetes, heart disease, and cancer (Chinnery & Gomez-Duran, 2018; Marom *et al*, 2017; Wei *et al*, 2017). We are certainly tantalized by the prospect that pathogenic variants illuminated by our approach might impinge upon human lifespan.

## METHODOLOGY

### Mitochondrial DNA sequence acquisition and conservation analysis

Mammalian mtDNA sequences were retrieved from the National Center for Biotechnology Information database of organelle genomes (https://www.ncbi.nlm.nih.gov/genome/browse#!/organelles/ on September 26, 2019). These 1184 mammalian mtDNA genomes were aligned using MAFFT on the ‘auto’ setting (Katoh & Standley, 2013). Four sequences that were egregiously misaligned were removed, and MAFFT alignment with the ‘auto’ setting was carried out again. Individual gene sequences were extracted from these alignments, based upon the annotated human mtDNA (NC_012920.1). After gap removal, translation of protein coding genes was performed using the vertebrate mitochondrial codon table in AliView (Larsson, 2014). MAFFT alignments of each gene product were performed using the G-INS-i iterative refinement method, then individual concatenates for each species were generated from protein coding sequences, tRNAs, and rRNAs. Duplicates were removed from the protein, tRNA, and rRNA concatenates using seqkit v0.10.2 (Shen *et al*, 2016).

Maximum likelihood trees for each molecule class concatenate were built using FastTreeMP (Price *et al*, 2010) with four subtree-prune-regraft moves, two rounds of branch length optimization, slow nearest neighbor interchange, and by use of a generalized time-reversible model. Next, ancestral prediction was carried out using the PAGAN package (Löytynoja *et al*, 2012), with concatenated alignments and phylogenetic trees used as input. The PAGAN output was then analyzed using “binary-table-by-edges-v2.2” and “add-convention-to-binarytable-v1.1.py” (https://github.com/corydunnlab/hummingbird) (Dunn *et al*, 2020) to allow for a sum of substitutions at alignment positions encoded by human mtDNA. All fluctuating edges were extracted using “report-on-F-values-v1.1” (https://github.com/corydunnlab/hummingbird). Correspondence files for the human mtDNA (NC_012920.1) convention and alignment positions were generated using “extract-correspondence-for-merged-alignment-v.1.1.py”, followed by the construction of the look-up tables of amino acid direct substitution and presence using “direct-subst-lookup-table-proteins-v.1.1.py” and “AA-presence-lookup-table-v.1.1.py”, respectively (https://github.com/corydunnlab/Edge_mapping_conservation). For IIDS determination, only edges with standard characters at both connected nodes were considered. For TSS and ISS calculation, the PAGAN output files for tRNAs and rRNAs were, before inferring a substitution, processed so that each ambiguous IUPAC nucleotide code was replaced by one of the possible standard nucleotides represented by that IUPAC code within the entire tree at that given position. For positions that contained only N’s and gaps, N’s were replaced by gaps. Replacement of ambiguous characters with standard characters was carried out using using script “replace-nonstandard.py” (https://github.com/corydunnlab/Edge_mapping_conservation), and the standard nucleotide was picked at random from available standard characters at a given alignment position using the random.choice() function of the Python random module. Gap percentage for each position was calculated using trimAI v1.4.rev22 (Capella-Gutiérrez *et al*, 2009) on alignments removed of predicted sequences for internal nodes. Shannon entropy was calculated using the ‘Shannon’ script (https://gist.github.com/jrjhealey/), also on alignments removed of predicted sequences for internal nodes. Custom scripts were used to merge various data tables. Software generated within the context of this study was written using Python 2.7.

Statistical testing and graph production were performed using Prism 8.4.3 (https://www.graphpad.com).

### mtDNA variant database utilization

MITOMAP data used in this study (Lott *et al*, 2013) were downloaded on October 1, 2019. HelixMTdb data used in this study (Bolze *et al*, 2019) were downloaded on October 15, 2019. Only substitutions were analyzed within the context of this study; insertions and deletions were not analyzed.

### Support vector machine classification

Our SVM classifier (Cortes & Vapnik, 1995) was developed using the R-language (https://www.R-project.org/) package e1071 (Meyer *et al*, 2014), for which the implementation is based on libsvm (Chang & Lin, 2011). A supervised binary classification model was built using positive and negative training sets. Each negative training set included those alleles with the highest number of homoplasmic samples in HelixMTdb and also annotated as a polymorphism within MITOMAP. For proteins, the negative training sets consisted of 50 mtDNA substitutions (encoding 51 protein variants) from the reference sequence. For tRNAs, the negative training sets also consisted of 50 substitutions. Positive training sets consisted of variants confirmed as pathogenic in MITOMAP on the date the database was queried. The positive training set for mtDNA-encoded proteins consisted of 39 mtDNA substitutions (encoding 40 protein variants), and the positive training set for mtDNA-encoded tRNAs consisted of 39 substitutions. Predictions were made for protein and tRNA sites gapped at 1.5% or less within our alignments. Decision values are related to, but do not directly represent, scalar distance from variant points to SVM margins (Sanz *et al*, 2018).

The best cost and gamma parameters for the classification model were selected based on majority voting of 100-searches of 2^(−8:8) grid surface, using ‘tune.svm’ function of e1071 [kernel = ‘radial’, type = ‘C-classification’, scale = TRUE, tunecontrol = tune.control (cross = 5, nrepeat = 10)]. The classification model was subsequently built and trained using the optimized [gamma, cost] parameters of [0.25, 0.0625] for proteins (Figure S9A) and [0.0078125, 256] for tRNAs (Figure S9B), and subsequently used to predict the pathogenicity of the training and test set variants. Features were scaled internally during training and when applying the model to predict the test set. Predictions for the ROC curve were collected using ‘mining’ function of the rminer package (Cortez, 2015), with the optimized parameters during 10 runs of 5-fold cross-validation [model=“ksvm”, task = “prob”, method = c(“kfold”, 5), Runs = 10]. Feature importance was measured using the ‘Importance’ function from rminer (Cortez & Embrechts, 2013), which was run 10 times (method = “DSA”).

Due to the relatively limited number of variants confirmed to be pathogenic or benign that one might include in a balanced training set, we also separately evaluated the robustness of our classifier using multiple runs of k-fold cross-validation with varying values of k. During these experiments, the complete list of our training set was divided into k parts, the classifier was trained on the (k-1) parts and the remaining part was used to evaluate the classifier performance. The k-fold cross-validation runs were performed with the optimized cost and gamma parameters given above. For both protein (Figure S9C) and tRNA (Figure S9D) training sets, 2-fold, 3.4-fold and 5-fold cross-validation was run 10 times, corresponding to setting 50%, 30% and 20%, respectively, of the training set aside for testing, then training the classifier with the remaining variants. In each run, mean performance metrics were collected by the ‘mining’ function over 10 repeats of the respective k-fold cross-validation [model = “ksvm”, task = “prob”, method = c(“kfold”, *variable*), runs = 10). Predictions for the set reserved for testing were then used to calculate performance metrics. The performance metrics MCC, Accuracy (ACC), True Positive Rate (TPR), True Negative Rate (TNR) and Area Under the ROC Curve (AUC) were calculated using ‘mmetric’ of the rminer package.

### Comparison of selected, alternative prediction methods with MitoCAP

Pathogenicity predictions for our training and test set variants were compared to predictions made by PolyPhen-2 (Adzhubei *et al*, 2013), PROVEAN (Choi *et al*, 2012), Panther-PSEP (Tang & Thomas, 2016b), Mitoclass (Martín-Navarro *et al*, 2017) and MitImpact (Castellana *et al*, 2015). PolyPhen-2 predictions were retrieved from its web server in batch query mode with the HimDiv classifier and default parameters. Similarly, PROVEAN predictions were retrieved from the web server for ‘PROVEAN Protein Batch’ tool for human proteins. For Panther-PSEP, variant queries for each of the 13 mitochondria-encoded proteins were generated and submitted, along with the protein sequence, to the web server. Mitoclass predictions were retrieved from the supplemental data of its publication (Martín-Navarro *et al*, 2017). For the meta-classifier employing its own SVM, MitImpact, the latest predictions (MitImpact 3D 3.0.5) were used. For comparison with PON-mt-tRNA (Niroula & Vihinen, 2016), tested variants were uploaded to the associated web server, and output was processed according to the mtDNA strand upon which the tRNA is encoded. The most recent MitoTIP (Sonney *et al*, 2017) predictions were published on April 27, 2020 and were downloaded directly from the MITOMAP server.

For PolyPhen-2 and Panther-PSEP, predictions ‘possibly damaging’ and ‘probably damaging’ were considered as ‘pathogenic’. For MitoTIP predictions, ‘likely pathogenic’ and ‘possibly pathogenic’ were collapsed to ‘pathogenic’ during comparisons, while ‘possibly benign’ and ‘likely benign’ were reduced to ‘benign’. Similarly, for PON-mt-tRNA annotations, ‘pathogenic’ and ‘likely pathogenic’ were reduced to ‘pathogenic’, while ‘likely neutral’ and ‘neutral’ annotations were considered as ‘benign’. Since MitImpact is a meta-classifier working on mtDNA sequence, it can produce two different pathogenicity predictions for two nucleotide changes leading to the same amino acid change. Such instances, where the same amino acid change is labeled both as ‘pathogenic’ and ‘neutral’ by MitImpact, were classified as ‘contradictory’ and not further processed during analyses.

## ACKNOWLEDGEMENTS

C.D.D. is funded by a ERC Starter Grant (RevMito 637649), by the Sigrid Jusélius Foundation, and by the Academy of Finland (331556). V.O.P. is supported by the Academy of Finland (289737, 314672, and 330255), U.S. National Institute of General Medical Sciences (1R01GM132649-01), the Sigrid Jusélius Foundation, and by the Runar Bäckström Foundation. We thank Vivek Sharma, Tamara Somborac, Svetlana Konovalova, and Gülayse Ince Dunn for helpful manuscript comments.

## AUTHOR CONTRIBUTIONS

B.A.A. developed software, analyzed data, and edited the manuscript. P.O.C. and V.O.P. analyzed data and edited the manuscript. C.D.D. conceived of the classification approach, supervised the project, analyzed data, prepared figures, and wrote the manuscript.

## SUPPLEMENTAL TABLE LEGENDS

**Table S1: SVM output analyzing protein-encoding variants found in the MITOMAP and HelixMTdb by use of mammalian sequence data**. Details of SVM analyses and features are found within the methodology section.

**Table S2: SVM output analyzing tRNA-encoding variants found in the MITOMAP and HelixMTdb by use of mammalian sequence data**. Details of SVM analyses and features are found within the methodology section.

---

**Figure S1:**
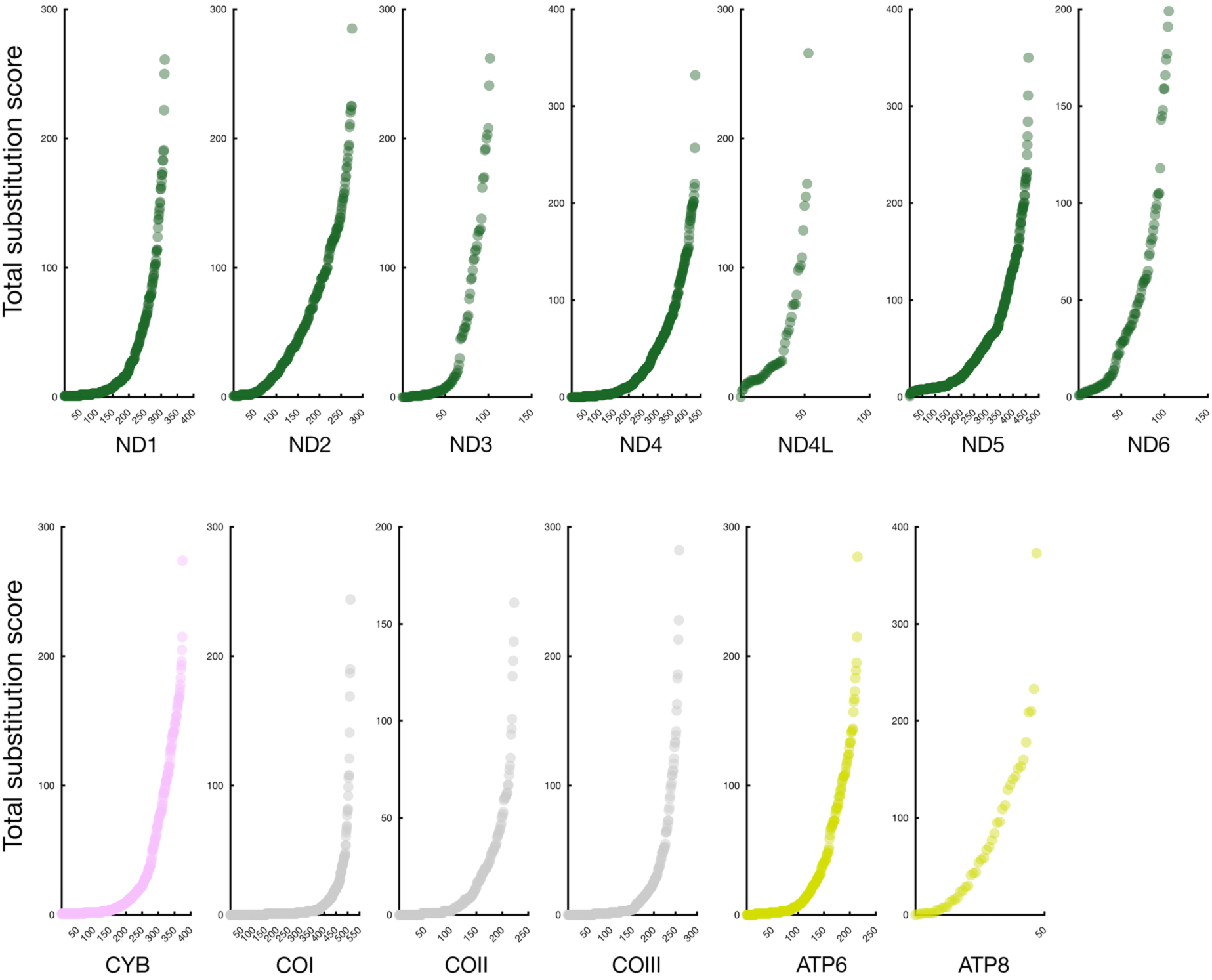
Distributions associated with site TSS values demonstrate that the majority of mtDNA-encoded protein sites are under selection and that site saturation is minimal or absent. Site TSSs, also provided in Figure 2A, are plotted in increasing order for each protein.

**Figure S2:**
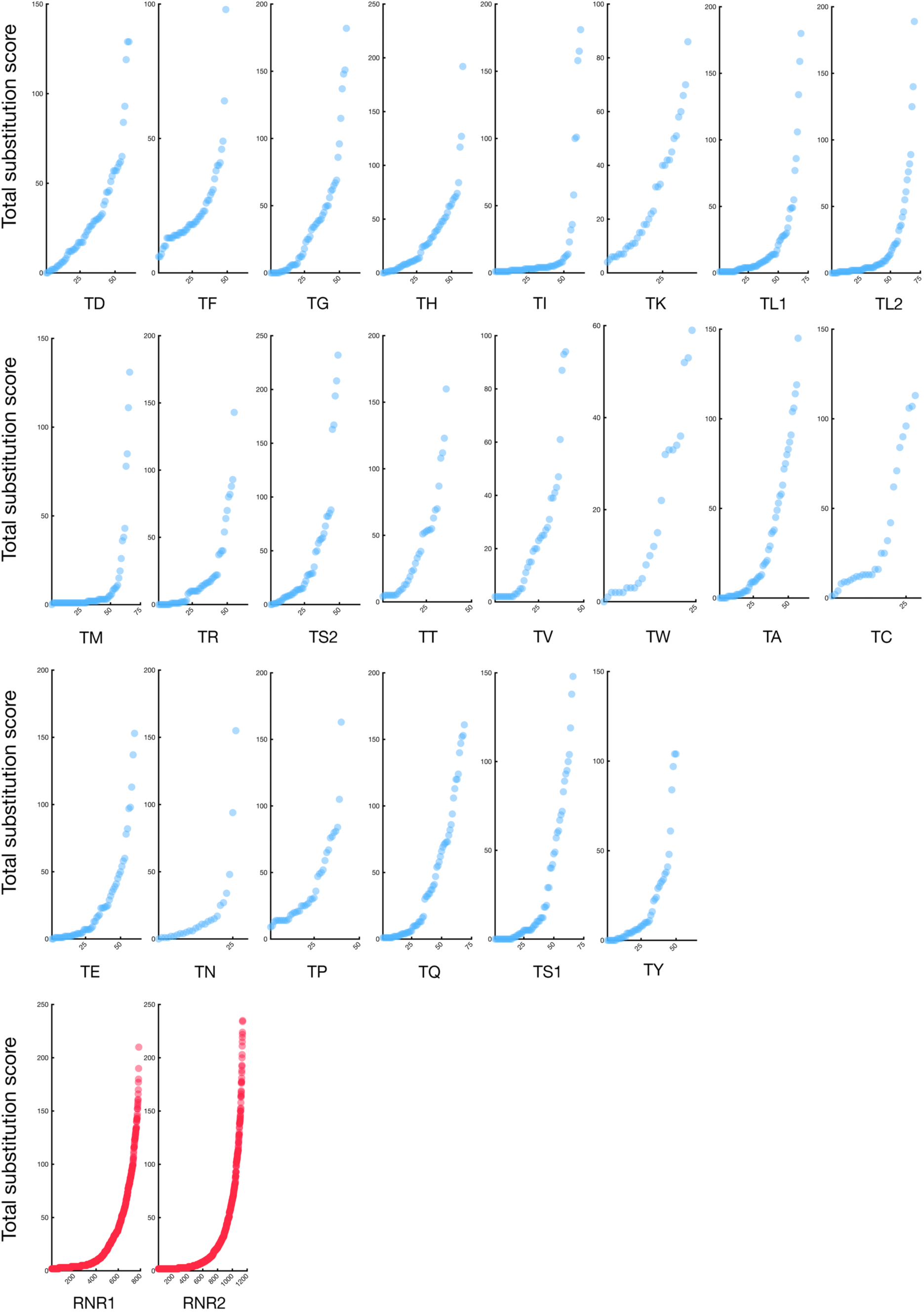
Distributions associated with site TSS values demonstrate that the majority of mtDNA-encoded RNA sites are under selection and that site saturation is minimal or absent. Plots are generated as in Figure S1, except tRNA and rRNA TSS distributions are shown.

**Figure S3:**
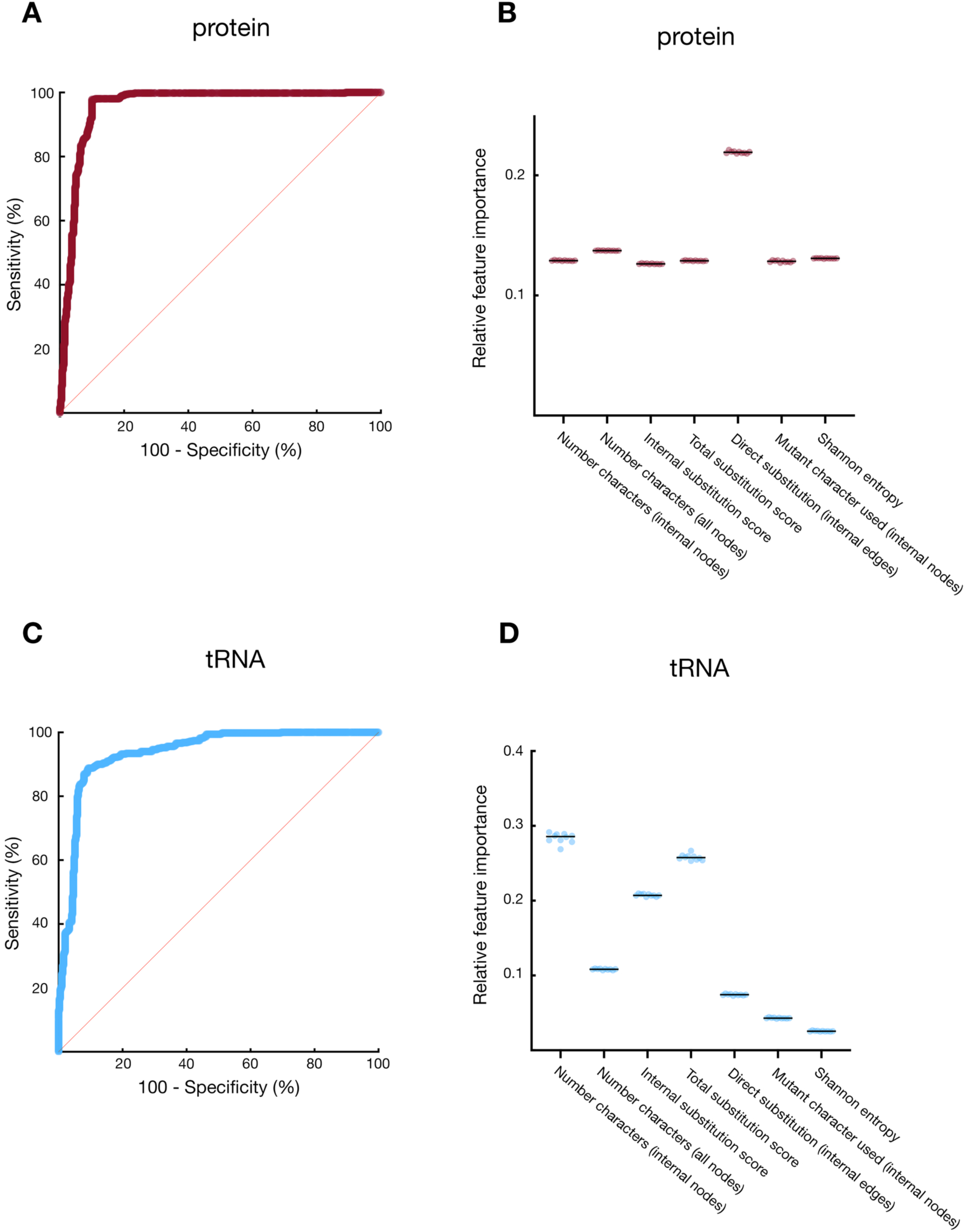
Feature importance and receiver operating characteristic curves for MitoCAP classification of protein and tRNA. (A) The ROC curve demonstrates very high sensitivity is achieved with no substantial loss in specificity for MitoCAP mammalian protein predictions. The true positive rate (sensitivity) of the predictive model is plotted against the false positive rate (1 - specificity) at various decision value thresholds. (B) Feature importance for prediction of protein-coding substitutions is calculated from the increase of the model’s prediction error after varying the values for each feature. (C) as in (A), except the ROC curve associated with tRNA substitution predictions is shown. (D) as in (B), except feature importance for tRNA variant predictions is provided.

**Figure S4:**
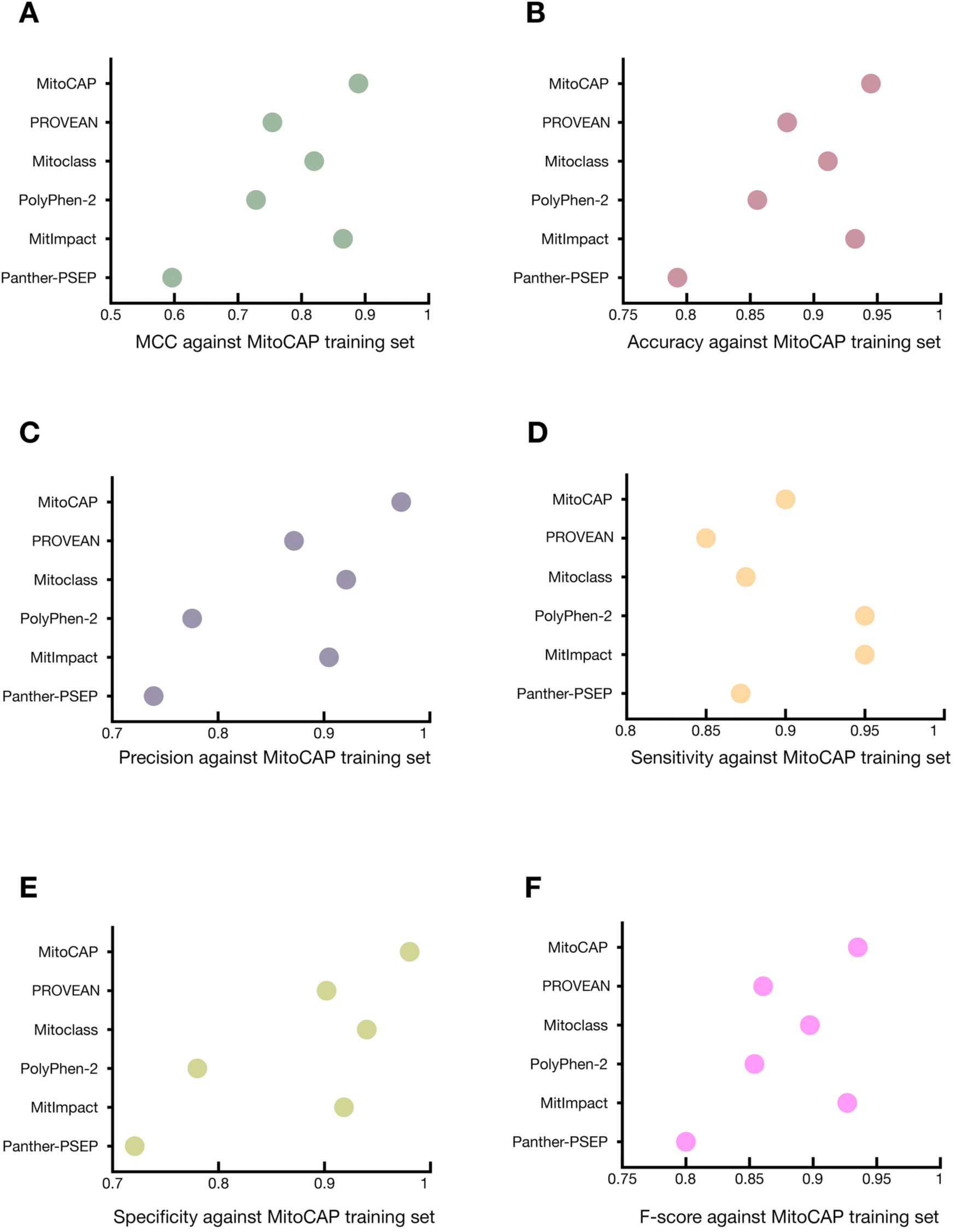
Metrics of classifier performance of MitoCAP and alternative methodologies using the MitoCAP training set for protein-coding substitutions. Plotted values obtained by the indicated classifiers include: (A) MCC, (B) accuracy, (C) precision, (D) sensitivity, (E) specificity, (F) F-score.

**Figure S5:**
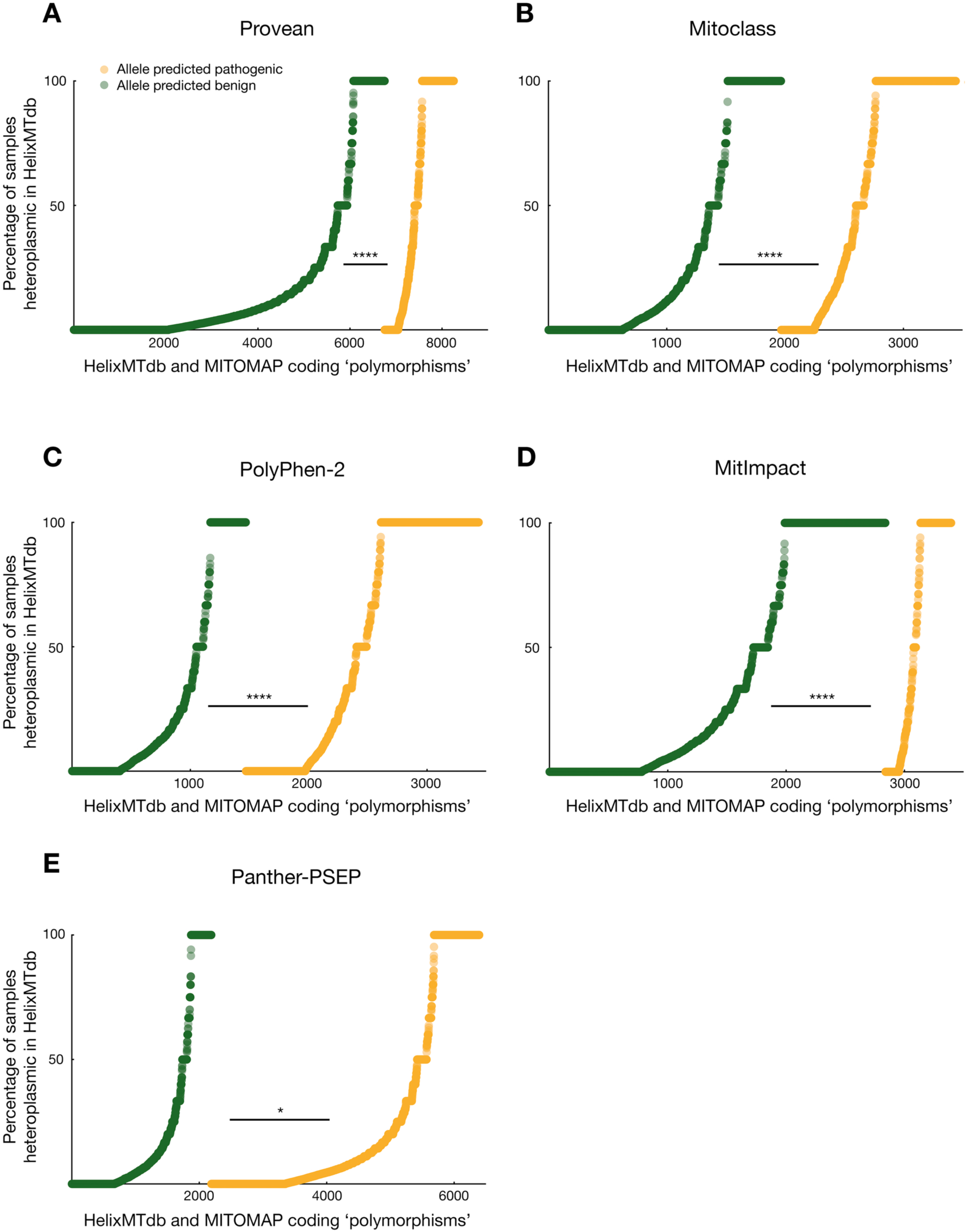
Separation of heteroplasmy frequency distributions for protein-coding substitutions by alternative classifiers. Distributions obtained using the indicated classifiers are plotted as in Figure 4B for (A) PROVEAN (B) Mitoclass (C) PolyPhen-2 (D) Mitlmpact (E) Panther-PSEP classifiers.

**Figure S6:**
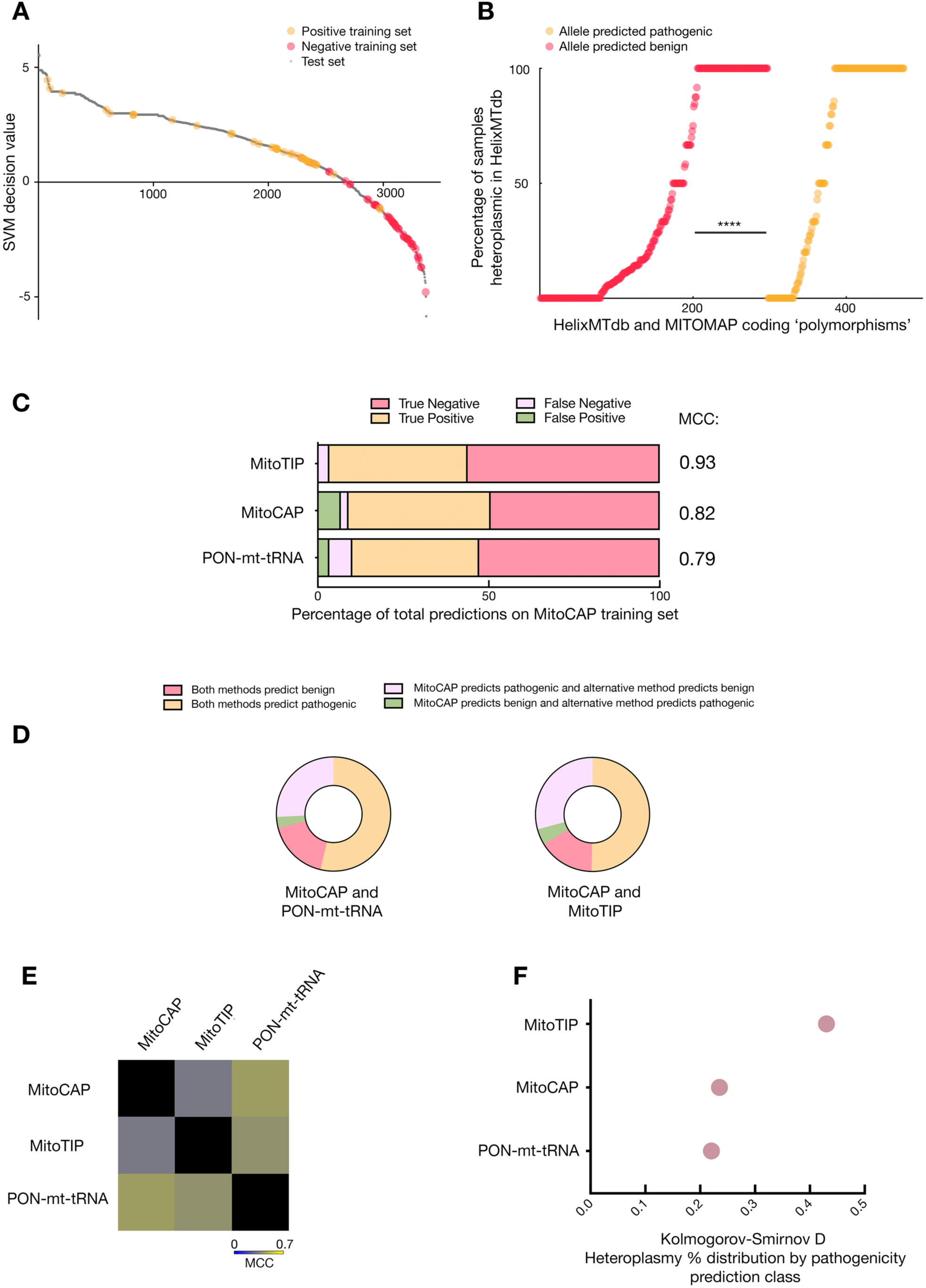
Prediction of mtDNA-encoded tRNA variants by the MitoCAP classifier. (A) as in Figure 4A, except that tRNA variants are plotted. (B) as in Figure 4B, except that tRNA variants are plotted. (C) as in Figure 4C, except that the outcome of other methods is compared to that of MitoCAP on the MitoCAP tRNA training set. (D-E) as in Figure 4D and Figure 4E, except that comparisons are made between the outcome of tRNA prediction models. (F) as in Figure 4F, except that distributions associated with heteroplasmy frequency and pathogenicity classification are made using tRNA substitution predictions.

**Figure S7:**
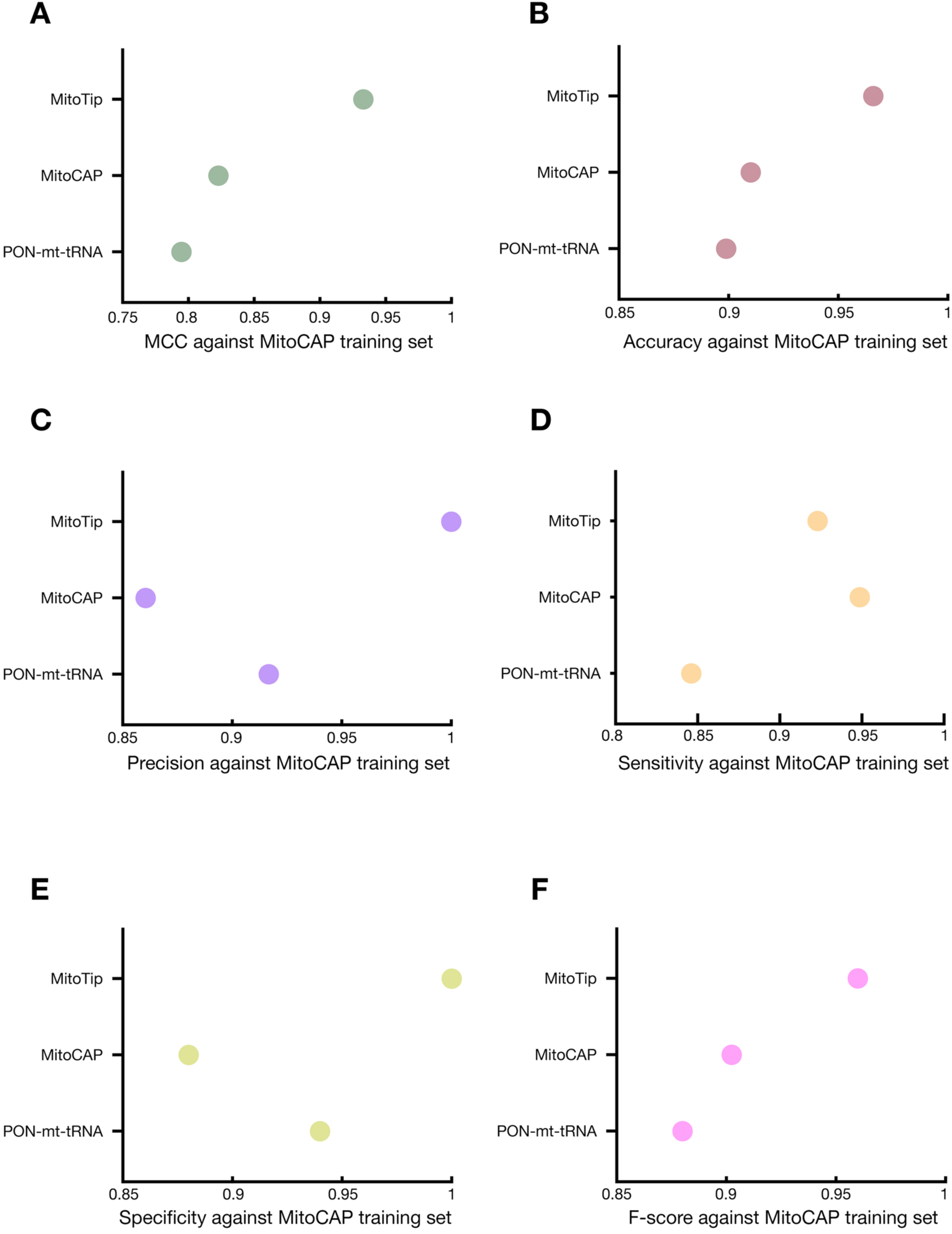
Metrics of classifier performance of MitoCAP and alternative methodologies on the MitoCAP training set for tRNA substitutions. Plotted values obtained by the indicated classifiers include: (A) MCC, (B) accuracy, (C) precision, (D) sensitivity, (E) specificity, (F) F-score.

**Figure S8:**
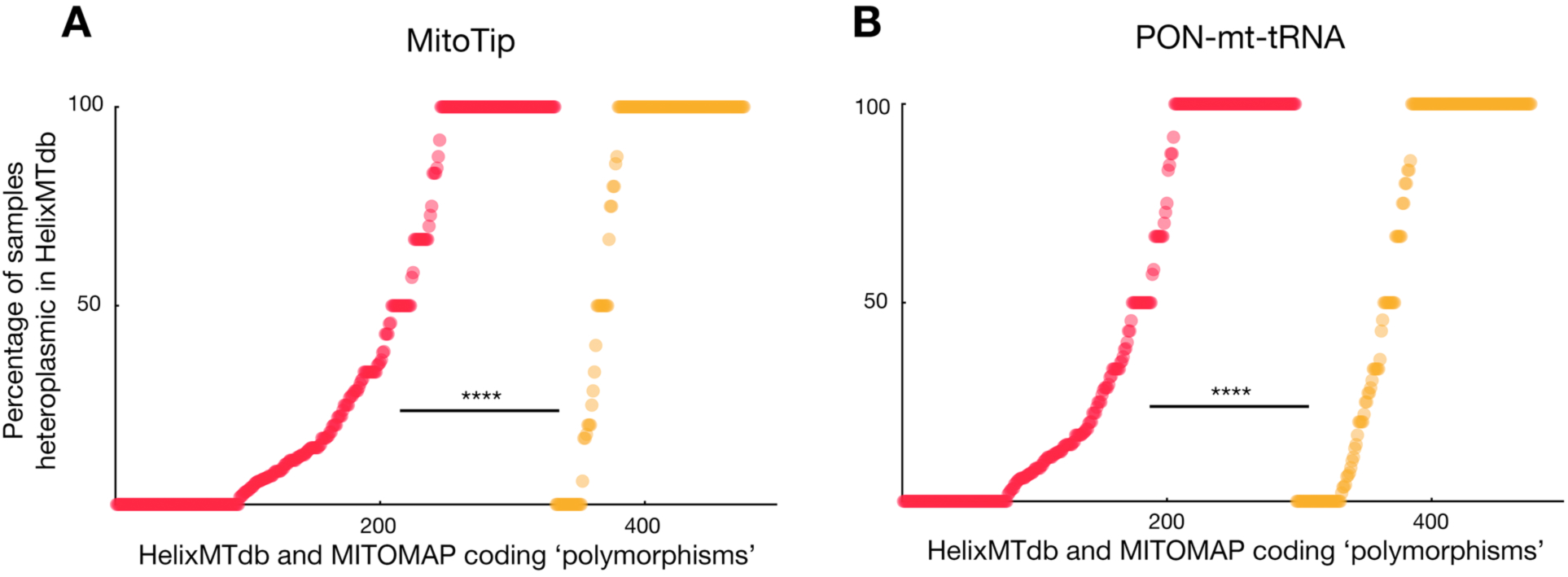
Separation of heteroplasmy frequency distributions for tRNA substitutions by alternative classifiers. Distributions obtained using the indicated classifiers are plotted as in Figure S6B for (A) MitoTIP and (B) PON-mt-tRNA classifiers.

**Figure S9:**
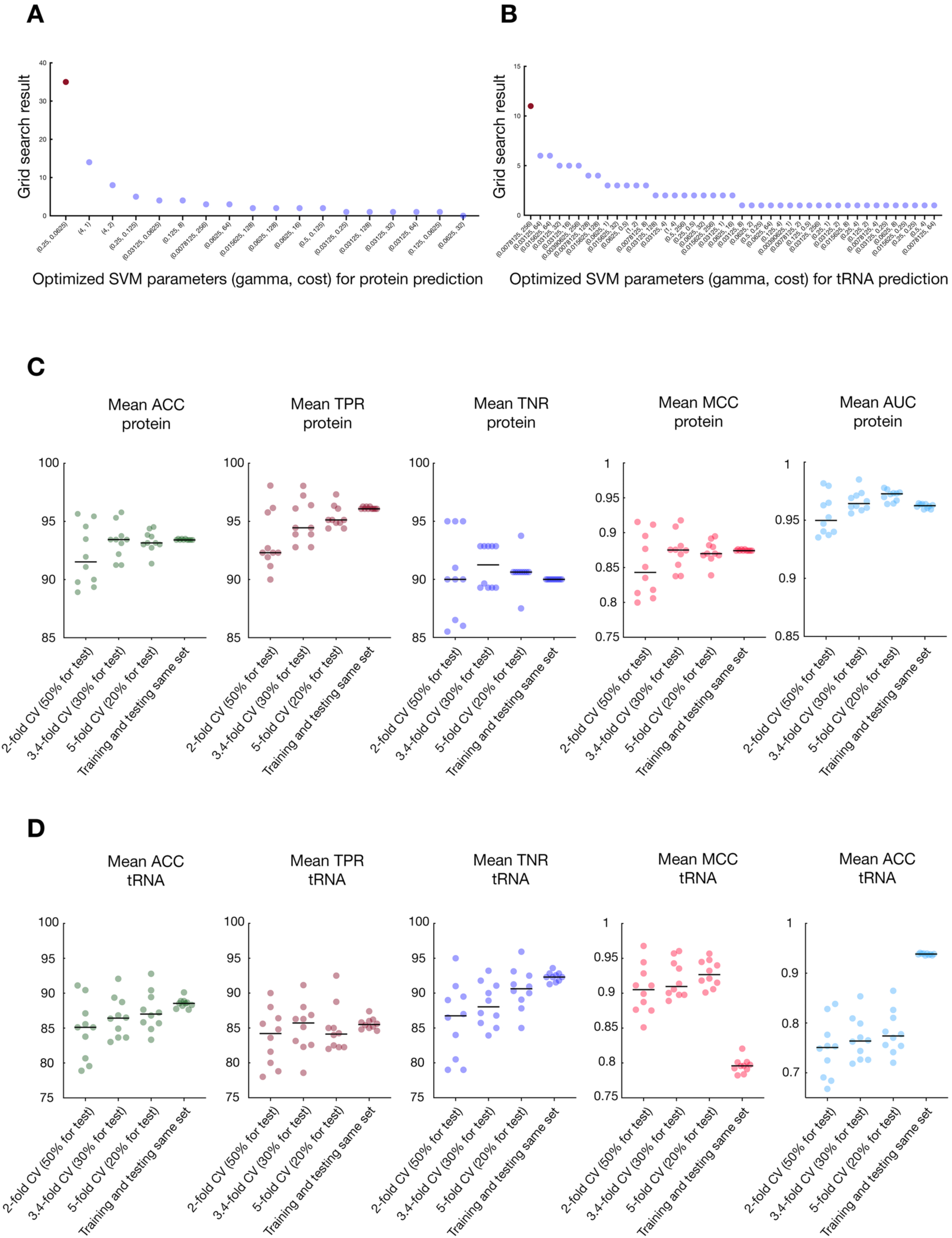
Optimization and validation of parameters used to develop the MitoCAP support vector machine classification model. (A) Optimal gamma and cost values for SVM model development were obtained before building the classifier. One hundred grid searches for optimal gamma and cost parameters were performed using the protein-coding substitution training dataset, and the values encountered the majority of the searches (red dot) were used during development of the classification model. (B) as for (A), but using the tRNA substitution training set. (C-D) Exploration of training set sufficiency demonstrates model stabilization using the full training sets. (C) k-fold cross-validation analyses were performed on the protein-coding variant training set to determine how robust classification might be to training set limitation. The mean values resulting from each instance of ten cross-validation tests with the indicated k-fold setting are plotted, and the bar indicates the median of these mean values. Mean accuracy (ACC), true positive rate (TPR), true negative rate (TNR), Matthews correlation coefficient (MCC), and area under the receiver operating characteristic curve (AUG) are plotted. (D) as in (C), except tRNA training sets were tested.

